# Contributions of the Dachsous intracellular domain to Dachsous-Fat signaling

**DOI:** 10.1101/2024.04.03.587940

**Authors:** Bipin Kumar Tripathi, Kenneth D. Irvine

## Abstract

The protocadherins Fat and Dachsous regulate organ growth, shape, patterning, and planar cell polarity. Although Dachsous and Fat have been described as ligand and receptor, respectively, in a signal transduction pathway, there is also evidence for bidirectional signaling. Here we assess signaling downstream of Dachsous through analysis of its intracellular domain. Genomic deletions of conserved sequences within *dachsous* identified regions of the intracellular domain required for normal development. Deletion of the A motif increased Dachsous protein levels and decreased wing size. Deletion of the D motif decreased Dachsous levels at cell membranes, increased wing size, and disrupted wing, leg and hindgut patterning and planar cell polarity. Co-immunoprecipitation experiments established that the D motif is necessary and sufficient for association of Dachsous with four key partners: Lowfat, Dachs, Spiny-legs, and MyoID. Subdivision of the D motif identified distinct regions that are preferentially responsible for association with Lft versus Dachs. Our results identify motifs that are essential for Dachsous function and are consistent with the hypothesis that the key function of Dachsous is regulation of Fat.

## Introduction

The Dachsous-Fat signaling pathway polarizes cells to regulate tissue polarity, growth, and patterning (reviewed by Fulford and McNeill, 2020; Gridnev and Misra, 2022; Strutt and Strutt, 2021). It is initiated by Dachsous (Ds) and Fat, two large cadherin proteins that bind each other through their extracellular domains (Clark et al., 1995; Ma et al., 2003; Mahoney et al., 1991; Matakatsu and Blair, 2004; Simon et al., 2010; Strutt and Strutt, 2002). Ds and Fat function in multiple organs throughout *Drosophila* development (reviewed by Fulford and McNeill, 2020; Gridnev and Misra, 2022; Strutt and Strutt, 2021). They are conserved in vertebrates, where they also function in multiple organs and have been linked to congenital diseases (Alders et al., 2014; Cappello et al., 2013; Durst et al., 2015; Mao et al., 2011a; Saburi et al., 2008; Zakaria et al., 2014). The *Drosophila* wing has been a key organ used to dissect Ds-Fat signaling. *ds* mutants have enlarged, rounder wings, abnormalities in wing patterning, and abnormal wing hair planar cell polarity (PCP) (Adler et al., 1998; Baena-Lopez et al., 2005; Clark et al., 1995). All of these phenotypes are shared by *fat* loss of function alleles or RNAi knockdown, but *ds* phenotypes are generally weaker than *fat* phenotypes, possibly because Fat can have some activity even in the absence of Ds (Bryant et al., 1988; Clark et al., 1995; Matakatsu and Blair, 2006). Binding between Ds and Fat is modulated by Four-jointed (Fj), a Golgi-localized kinase that phosphorylates their cadherin domains (Brittle et al., 2010; Ishikawa et al., 2008; Simon et al., 2010; Strutt and Strutt, 2002). Ds, Fj, and in some cases Fat are expressed in gradients across tissues where they function, and their differential expression and binding interactions leads to polarized membrane localization of Ds and Fat (Ambegaonkar et al., 2012; Brittle et al., 2012; Cho and Irvine, 2004; Clark et al., 1995; Ma et al., 2003; Matakatsu and Blair, 2004; Strutt and Strutt, 2002; Villano and Katz, 1995).

Ds-Fat regulate gene expression through the Hippo signaling pathway, and in this context Ds and Fat have been described as ligand and receptor, respectively (Bennett and Harvey, 2006; Cho et al., 2006; Mao et al., 2006; Matakatsu and Blair, 2006; Rogulja et al., 2008; Silva et al., 2006; Willecke et al., 2006; Willecke et al., 2008). However, other observations suggest that Ds-Fat signaling should be considered as bidirectional, implying that they function as both ligand and receptor for each other, and PCP signaling is inherently bidirectional (Casal et al., 2006; Degoutin et al., 2013; Matakatsu and Blair, 2012; Zecca and Struhl, 2010). Moreover, the observation that elimination of both Ds and Fat results in stronger phenotypes than elimination of Fat alone implies that Ds has functions beyond regulation of Fat (Matakatsu and Blair, 2006). Extensive studies of the Fat intracellular domain (ICD) have provided insights into how it mediates downstream signal transduction and supported its classification as a signal-transducing receptor (Fulford et al., 2023; Matakatsu and Blair, 2012; Pan et al., 2013; Zhao et al., 2013). However, comparable studies have not previously been described for Ds. Here, we remedy this by describing functional and biochemical analysis of conserved sequence motifs within the Ds ICD.

Several proteins that act at different points within Ds-Fat signaling have been identified. One factor that is highly conserved in vertebrates is Lowfat (Lft), which is required to maintain normal levels of both Fat and Ds in the developing *Drosophila* wing, and which can physically associate with the cytoplasmic domains of Fat and Ds (Mao et al., 2009). A key downstream factor mediating the influence of Ds-Fat signaling on both Hippo and PCP pathways in *Drosophila* is the atypical myosin Dachs (Ambegaonkar and Irvine, 2015; Ayukawa et al., 2014; Bosveld et al., 2012; Cho et al., 2006; Cho and Irvine, 2004; Mao et al., 2006; Mao et al., 2011b). Ds-Fat signaling regulates the levels of Dachs membrane localization to modulate Hippo signaling, and the polarity of Dachs membrane localization to modulate PCP. In the developing wing imaginal disc, Dachs is localized to the distal sides of cells, where it often co-localizes with Ds (Ambegaonkar et al., 2012; Brittle et al., 2012; Mao et al., 2006). Dachs is removed from the proximal sides of cells in Fat-dependent process, and mutation or knockdown of *fat* leads to Dachs around the entire cell circumference, while over-expression of Fat removes Dachs from cell membranes (Mao et al., 2006). The significance of Ds-Dachs co-localization, and of the ability of Ds and Dachs to physically associate, has remained unclear. Dachs, together with the co-dependent factor Dlish/Vamana, regulate Hippo signaling through effects on the levels and activity of the Hippo pathway kinase Warts, and the upstream Hippo pathway regulator Expanded (Cho et al., 2006; Feng and Irvine, 2007; Misra and Irvine, 2016; Vrabioiu and Struhl, 2015; Wang et al., 2019; Zhang et al., 2016). Ds-Fat polarize cells in part by regulating the Fz, or core, PCP pathway, and in part independently of Fz PCP signaling (reviewed in Strutt and Strutt, 2021). Crosstalk with Fz-PCP signaling is mediated through Spiny-legs (Sple), which is an isoform of Prickle (Pk) (Ambegaonkar and Irvine, 2015; Ayukawa et al., 2014; Gubb et al., 1999; Merkel et al., 2014; Olofsson et al., 2014). Dachs and Ds can each physically associate with Sple and contribute to polarized Sple localization within cells.

Here, we use a structure-function approach to investigate potential signal transduction downstream of Ds. Ds is a 379 kD protein including a large extracellular domain with 27 cadherin repeats and a small (436 aa) intracellular domain (Clark et al., 1995). We identified and deleted six different conserved motifs within the Ds ICD. Phenotypic analysis identified contributions of these motifs to Ds activity. Part of this can be ascribed to effects of these motifs on Ds protein levels and distribution, as deletion of the A motif increases Ds protein levels, while deletion of the D motif decreases Ds protein at cell membranes. The change in Ds protein in the absence of the D motif can be explained by our discovery that it associates with and is required for regulation of Ds by Lft. The D motif also associates with other factors important to Ds-Fat signaling, including Dachs, Sple and MyoID, and by making smaller deletions of the D motif the effects of Lft binding could be partially separated from other requirements for this motif.

## Results

### Identification and deletion of conserved sequence motifs in the Ds ICD

We assessed evolutionary conservation to guide identification of functionally important regions of the Ds ICD. Conserved amino acids identified by a sequence alignment tool (Clustal Omega) were clustered in six different regions (A through F), extending from near the transmembrane domain to the C-terminus (Figures 1B, Supplemental S1). Four of these motifs (B, D, E, and F) are conserved from insects to vertebrates, whereas two (A and C) are only conserved within insects.

**Figure 1.**
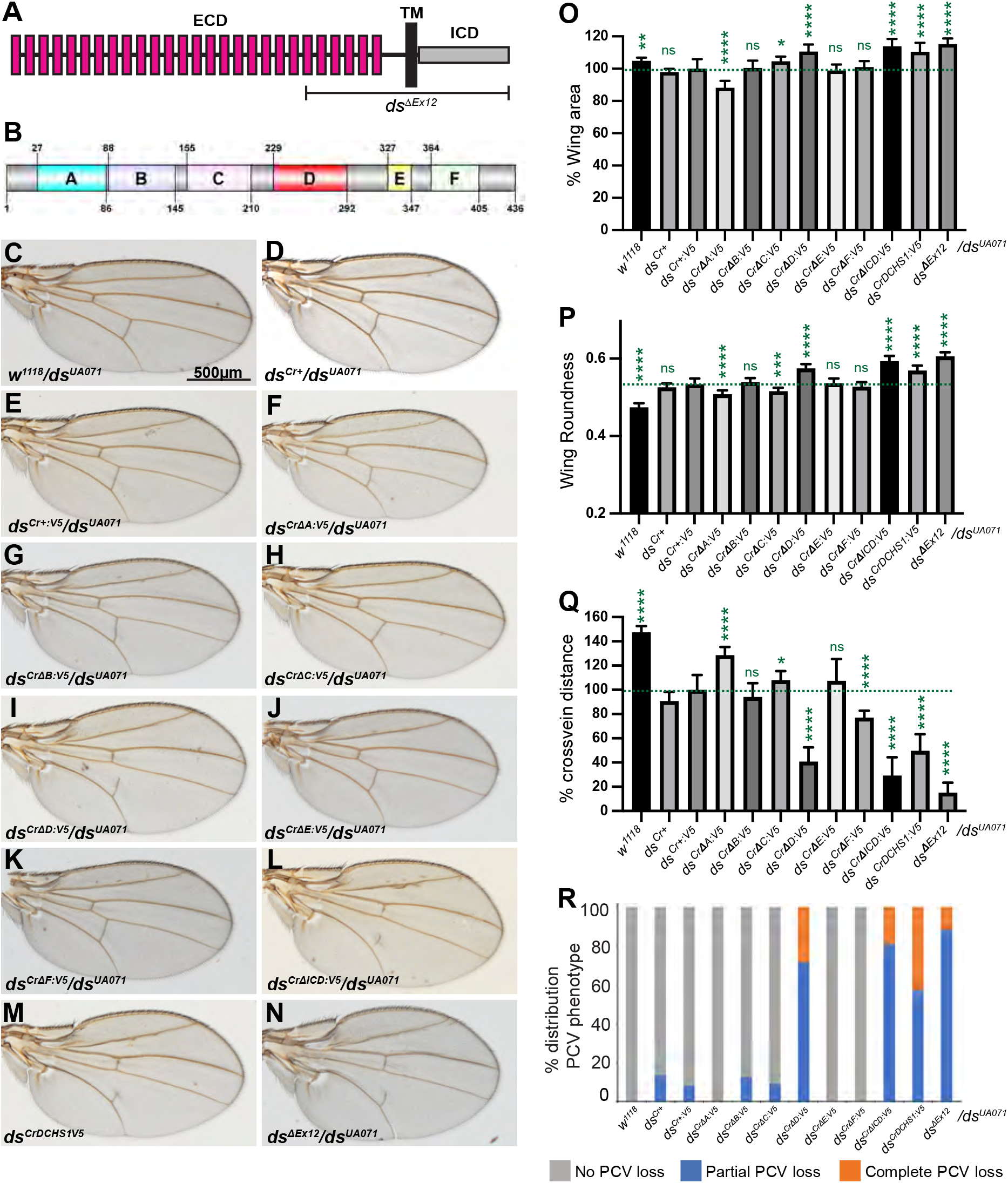
Adult wing phenotypes of Ds-ICD motif deletions. **(A)** Schematic of Ds, depicting cadherin repeats (red rectangles) in the extracellular domain (ECD), transmembrane domain (TM), and intracellular domain (ICD). The region affected by exon 12 deletion is indicated (*ds*^Δ*Ex12*^). **(B)** Schematic of the location of conserved sequence motifs within the Ds-ICD. **(C-N)** Adult male wings from *w^1118^/ds^UA071^*, n=26 *(C), ds^Cr+^/ds^UA071^*, n=15 *(D), ds^Cr+:V5^/ds^UA071^*, n=20 *(E), ds^Cr^*^Δ^*^A:V5^/ds^UA071^*, n=18 *(F), ds^Cr^*^Δ^*^B:V5^/ds^UA071^*, n=24 *(G), ds^Cr^*^Δ^*^C:V5^/ds^UA071^*, n=22 *(H), ds^Cr^*^Δ^*^D:V5^/ds^UA071^*, n=19 *(I), ds^Cr^*^Δ^*^E:V5^/ds^UA071^*, n=18 *(J), ds^Cr^*^Δ^*^F:V5^/ds^UA071^*, n=23 *(K), ds^Cr^*^Δ^*^ICD:V5^/ds^UA071^*, n=16 *(L), ds^CrDCHS1:V5^/ds^UA071^*, n=14 *(M), and ds*^Δ^*^Ex12^/ds^UA071^*, n=30 *(N)*. Scale bar = 500µm. **(O-R)** Histograms illustrating relative wing area (as compared to that in *ds^Cr+:V5^/ds^UA071^*) (O), wing roundness (P), relative cross-vein distance (Q), and PCV loss (R) in the indicated genotypes. Error bar indicates mean ± SD from the number of wings indicated above.

The ICD, transmembrane domain, and part of the extracellular domain are encoded by a single large exon of *ds* (exon 12) (Figures 1A, Supplemental S2A). To assess the functional significance of conserved motifs, we used a recombination-mediated cassette exchange (RMCE) strategy (Zhang et al., 2014), which enables efficient replacement through site-specific recombination of sequences flanked by attP sites. CRISPR-Cas9 mediated recombination was used to replace exon 12 with an RCME cassette including an RFP marker gene, flanked by attP sites in the upstream intron and 3’ UTR of *ds* (Supplemental Figure S2A). This replacement of exon 12 created a new mutant allele, *ds*^*ΔEx12*^, which behaves as a strong *ds* allele in genetic crosses, resulting in characteristic *ds* phenotypes including wing overgrowth, rounder wing shape, reduced cross-vein spacing, abnormal wing hair polarity, and partial loss of the posterior cross-vein (PCV) (Supplemental Figure S2). The RMCE cassette within *ds* was then replaced by site-specific recombination with either wild-type *ds* sequences, or with *ds* sequences in which one of the motifs A through F were deleted. We also created alleles in which the entire *ds* ICD was deleted (*ds*^*Cr*Δ*ICD:V5*^), and in which the entire ICD was replaced by the ICD from human DCHS1 (*ds^CrDCHS1:V5^*). All these also included a C-terminal V5 tag, connected by a flexible linker, so that the levels and distribution of the encoded Ds proteins could be examined.

### Identification of conserved motifs that contribute to Ds function in wings and legs

To assess the activity of each *ds* allele, we examined them in transheterozygous combinations over the previously described strong *ds* allele *ds^UA071^*. We examined characteristic *ds* wing phenotypes, including wing size, shape, cross-vein spacing, posterior cross-vein loss, and wing hair polarity. Unexpectedly, our allele in which the RMCE cassette was replaced by wildtype *ds* sequences, *ds^Cr+:V5^*, behaves as a weak *ds* allele. *ds^Cr+:V5^* wings are slightly smaller and rounder than wildtype (*w^1118^*) control wings (Figure 1C, E, O, and P), and the distance between the anterior and posterior cross-veins is reduced (Figure 1Q). Wing hair polarity is, however, normal (Figure 2A, E). We note that the influence of *ds* on wing size is complex, as wings in animals with strong *ds* alleles are larger than wild-type, whereas wings in animals with weak *ds* alleles are slightly smaller than wild-type (Clark et al., 1995).

**Figure 2.**
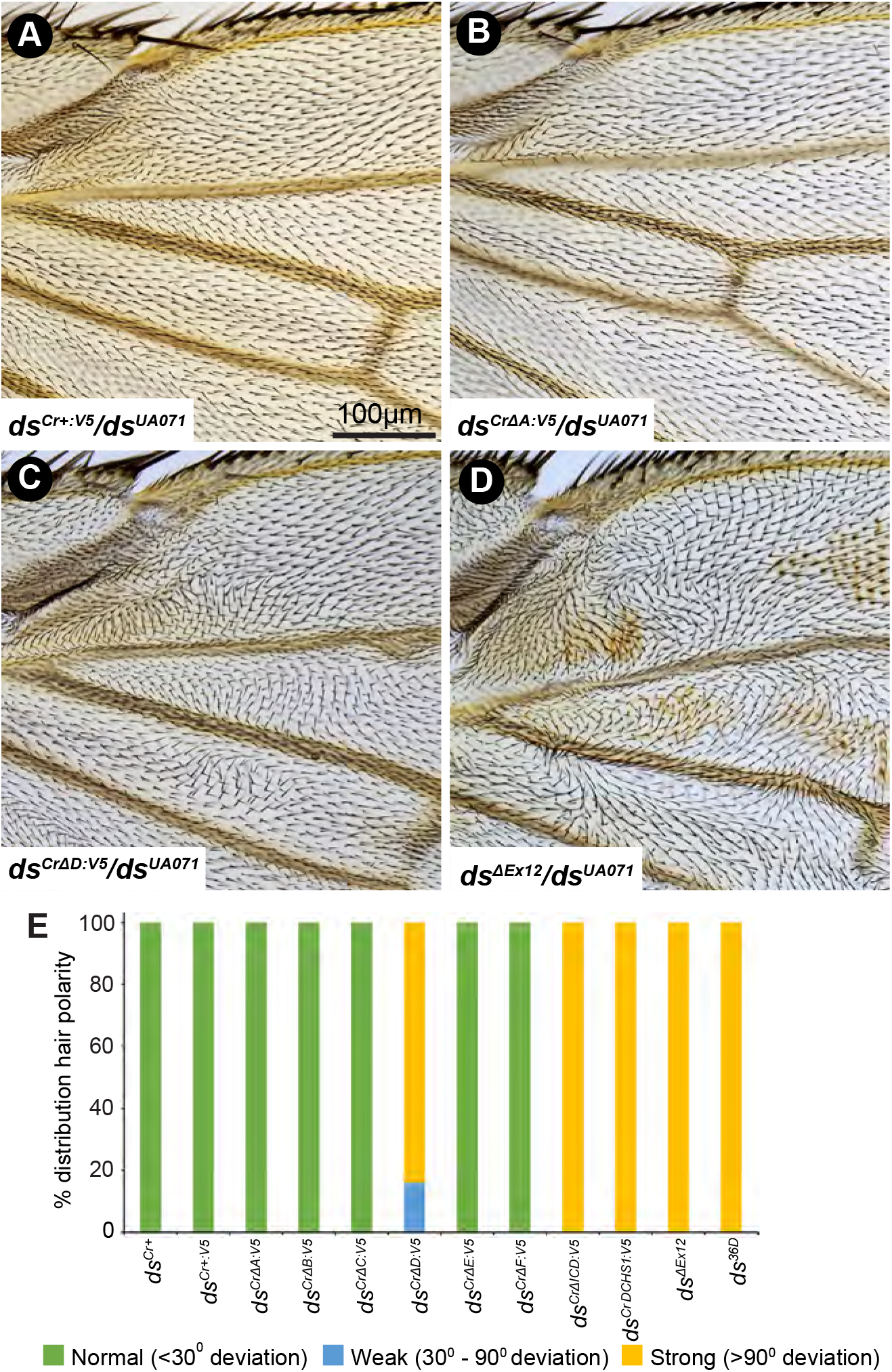
Wing hair PCP phenotypes of Ds-ICD motif deletions. **(A-D)** Proximal anterior wing of adult males from *ds^Cr+:V5^/ds^UA071^ (A), ds^Cr^*^*ΔA:V5*^*/ds^UA071^ (B), ds^Cr^*^*ΔD:V5*^*/ds^UA071^(C),* and *ds*^*ΔEx12*^*/ds^UA071^(D).* Scale bar = 100µm. **(E)** Histogram showing distribution of proximal wing hair PCP in different alleles transheterozygous over *ds^UA071^*based on analysis of 14 to 30 wings per genotype (as listed in Figure 1 legend). Wings were scored as normal if wing hairs deviate less than 30° from normal, weak if at least 10% of wing hairs deviate 30-90° from normal, and strong if at least 10% of wing hairs deviate >90° from normal.

To investigate whether the insertion of the V5 tag was interfering with Ds activity, we created an additional RMCE-mediated wildtype replacement allele without the V5 tag (*ds^Cr+^*). However, comparison of *ds^Cr+^* and *ds^Cr+:V5^* revealed that they have similar phenotypes, including mild reductions in wing growth and cross-vein spacing, together with slightly increased wing roundness (Figure 1D, O-Q). Sequencing of genomic DNA in *ds^Cr+:V5^* including all of exon 12 and an additional 1kb upstream and downstream did not reveal any alterations in the sequence other than the attP sequences in non-coding regions used to mediate RMCE. Thus, we conclude that the V5 tag does not affect Ds protein, but the inserted attP sequences may result in mild alterations of *ds* expression. As our goal was to compare the activities of different proteins, we continued with analysis of the different deletion alleles we had created, using *ds^Cr+:V5^* as our baseline for comparison to wild-type protein.

Three of the six alleles with deletions of conserved sequence motifs exhibited altered wing sizes as compared to *ds^Cr+:V5^*. *ds*^*CrΔA:V5*^ wings average 12 % smaller than control wings (Figure 1F, O), whereas *ds*^*Cr*Δ*C:V5*^ and *ds*^*Cr*Δ*D:V5*^ wings average 5% and 11%, respectively, larger than control wings (Figure 1H, I, O). Conversely, wings in *ds*^*Cr*Δ*B:V5*^, *ds*^*Cr*Δ*E:V5*^ or *ds*^*Cr*Δ*F:V5*^ flies do not differ significantly in size from controls (Figure 1G, J, K, O). In flies with deletion of the entire ICD, *ds*^*Cr*Δ*ICD:V5*^, wings were 14 % larger than the *ds^Cr+:V5^* wild type control (Figure 1L, O), similar to the 16% increase in wing size in *ds*^*ΔEx12*^ flies (Figure 1N, O). In flies expressing *ds* with the human DCHS1 ICD, wings were 10% larger than controls (Figure 1M, O).

The wing normally has an elongated shape, and both weak and strong *ds* alleles result in rounder wings (Clark et al., 1995). Wing roundness (measured as 4 ×[(Area)/π × (Major axis)^2^]) is significantly increased in *ds*^*Cr*Δ*D:V5*^ wings as compared to *ds^Cr+:V5^* controls, but slightly reduced in *ds*^*CrΔA:V5*^ and *ds*^*Cr*Δ*C:V5*^ wings (Figure 1P). Wing shape in *ds*^*Cr*Δ*B:V5*^, *ds*^*Cr*Δ*E:V5*^ and *ds*^*Cr*Δ*F:V5*^ does not differ significantly from controls (Figure 1P). *ds*^*ΔEx12*^, *ds*^*Cr*Δ*ICD:V5*^ and *ds^CrDCHS1:V5^* wings were all significantly rounder than control wings (Figure 1P).

Reduced distance between anterior and posterior cross-veins is a characteristic feature of mutations in many genes in the Ds-Fat pathway, including *ds*. Amongst the motif deletion alleles that we created, deletion of the D-motif, in *ds*^*Cr*Δ*D:V5*^, has the strongest effect, with cross-vein spacing reduced by 59% compared to that of control wings (Figure 1I, Q). Cross-vein spacing is also reduced by 23% in *ds*^*Cr*Δ*F:V5*^ (Figure 1K, Q). Cross-vein spacing is increased compared to control by deletion of the A motif (by 28%) or deletion of the C motif (by 8%) (Figure 1F, H, Q). No significant difference in cross-vein spacing compared to control is observed in wings expressing the *ds*^*Cr*Δ*B:V5*^ or *ds*^*Cr*Δ*E:V5*^ alleles (Figure 1G, J, Q). Deletion of the entire ICD, in *ds*^*Cr*Δ*ICD:V5*^, results in a 71% reduction in cross-vein distance compared to control wings, and cross-vein distance is reduced by 85% in *ds*^*ΔEx12*^ (Figure 1L, N, Q). In *ds^CrDCHS1:V5^* wings, cross-vein spacing was reduced by 50% (Figure 1M, Q).

*ds* mutant alleles also often exhibit a partial loss of the posterior cross vein (PCV). We categorized these phenotypes as: no PCV loss, partial PCV loss, or complete PCV loss. In *ds^Cr+:V5^* control flies, 92% have no PCV loss, and 8% have partial PCV loss (Figure 1R). Wings from *ds*^*Cr*Δ*B:V5*^ and *ds*^*Cr*Δ*C:V5*^ flies have PCV loss phenotypes similar to control flies (88% and 91% with no PCV loss, respectively). 100% of *ds*^*CrΔA:V5*^; *ds*^*Cr*Δ*E:V5*^ and *ds*^*Cr*Δ*F:V5*^ wings have no PCV loss (Figure 1R). Conversely, PCV loss was increased in *ds*^*Cr*Δ*D:V5*^ wings, with 29% having complete PCV loss and the remaining 71% with partial PCV loss. Strong PCV loss phenotypes were also observed in *ds*^*Cr*Δ*ICD:V5*^, *ds*^*CrDCHS1:V5*^, and *ds*^*ΔEx12*^ (Figure 1R).

Wing hairs are normally oriented from proximal to distal, but hair polarity is disrupted in PCP mutants. Ds-Fat pathway mutants have their strongest effects in proximal regions of the wing, and in adult wings from *ds* mutants misoriented hairs and swirling patterns can be readily observed in the anterior proximal wing region (Figure 2D). Control flies expressing wild-type Ds protein (*ds^Cr+:V5^*) or flies expressing *ds*^*CrΔA:V5*^ do not exhibit wing hair PCP phenotypes (Figure 2A, B, E). Flies expressing *ds*^*ΔEx12*^ *ds*^*Cr*Δ*ICD:V5*^, or *ds^CrDCHS1:V5^* have strong hair polarity defects (hairs deviated more than 90° from normal orientation) in 100 % of wings examined (Figures 2 D, E, Supplemental S3F, G). *ds*^*Cr*Δ*D:V5*^ also has consistent hair polarity defects in the proximal anterior wing, with 84% of wings examined showing strong polarity defects and 16% of wings showing weak polarity defects (hairs deviated between 30° and 90° from normal orientation) (Figure 2C, E). Wings from *ds*^*Cr*Δ*B:V5*^; *ds*^*Cr*Δ*C:V5*^; *ds*^*Cr*Δ*E:V5*^ and *ds*^*Cr*Δ*F:V5*^ flies have no evident wing hair polarity phenotypes (Figures 2E, Supplemental S3B-E).

We also analyzed leg phenotypes. In *ds* mutants, legs are shorter and wider than wild-type legs and the number of tarsal segments is reduced (Supplemental Figure S3S, T) (Clark et al., 1995). A reduced number of tarsal segments was observed in *ds*^*Cr*Δ*D:V5*^ flies, as well as *ds*^*Cr*Δ*ICD:V5*^*, ds^CrDCHS1:V5^*, and *ds*^*ΔEx12*^ flies (Supplemental Figure S3M, P, Q, R). Conversely, *ds*^*CrΔA:V5*^; *ds*^*Cr*Δ*B:V5*^; *ds*^*Cr*Δ*C:V5*^; *ds*^*Cr*Δ*E:V5*^ and *ds*^*Cr*Δ*F:V5*^ lines all have 5 tarsal segments, as observed in control flies *ds^Cr+:V5^* and *w^1118^* (Supplemental Figure S3I-L, N, O).

### Influence of Ds-ICD deletions on Ds protein localization and levels

To investigate the basis for the influence of these alleles on Ds activity, we first examined the localization and levels of the Ds protein expressed by them in wing imaginal discs. Examination of Ds:V5 protein expressed by *ds^Cr+:V5^* revealed a protein distribution similar to that previously described for Ds, including a proximal to distal gradient with relatively high levels in the wing hinge, low levels in the proximal wing pouch, and barely detectable levels in the distal wing pouch (Figure 3). As for endogenous Ds, Ds:V5 protein is localized near apical cell junctions, and at higher magnification often appears somewhat punctate (Figure 3C).

**Figure 3.**
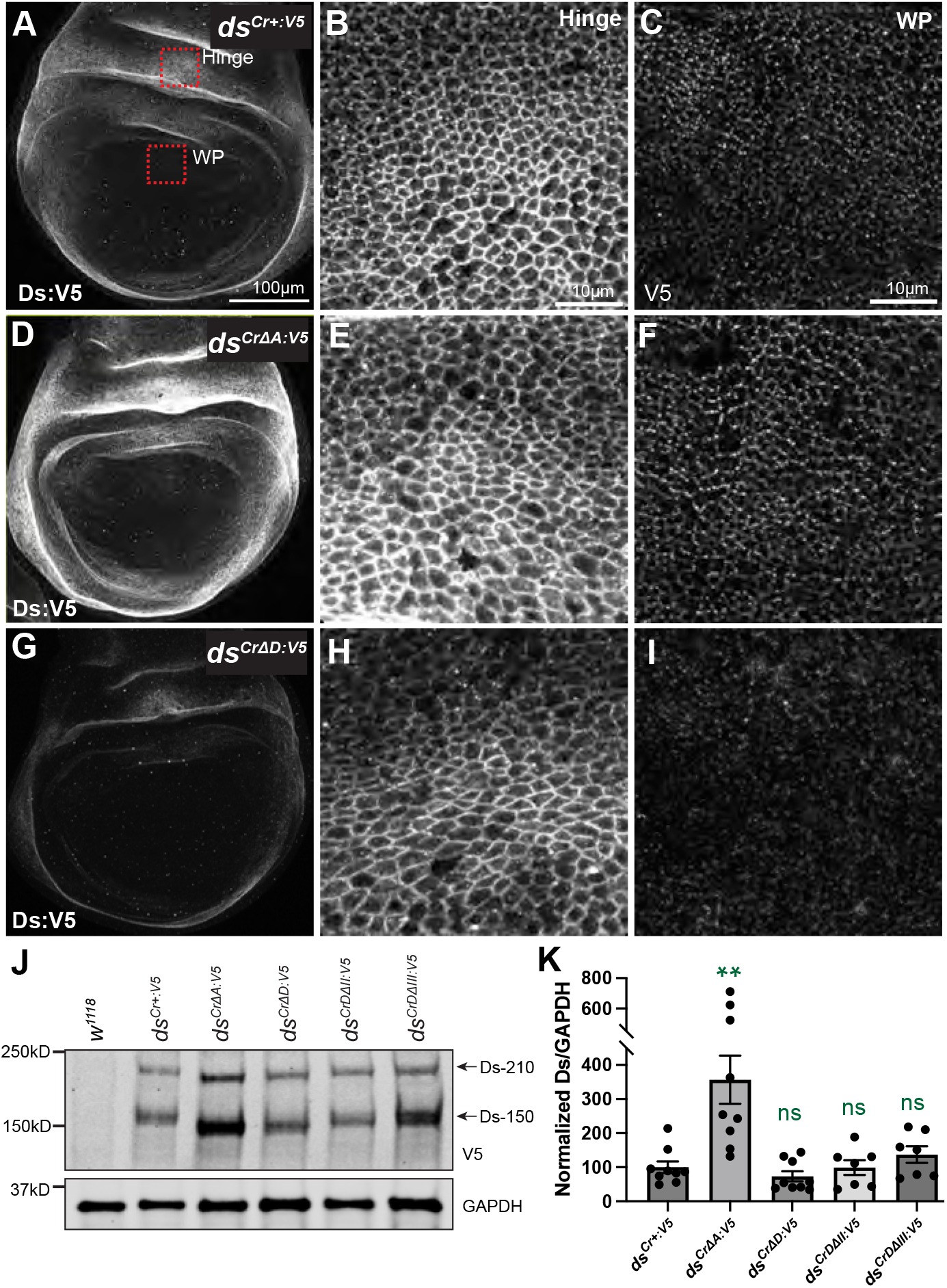
Localization and levels of Ds protein. Third-instar wing discs expressing Ds:V5 protein from homozygous *ds^Cr+:V5^* (A-C), *ds*^*CrΔA:V5*^ (D-F), and *ds*^*Cr*Δ*D:V5*^ (G-I). Red square boxes in (A) show approximate locations in hinge and wing pouch (WP) corresponding to the higher magnification panels at right. (**A,D,G**) Lower magnification images showing the entire wing pouch and hinge regions, Scale bar = 100 µm. **(B,E,H)** Images depicting part of the wing hinge region from homozygous *ds^Cr+:V5^* (B), *ds*^*Cr*Δ*A:V5*^(E), and *ds*^*Cr*Δ*D:V5*^(H). Scale bar = 10 µm. **(C,F,I)** Images depicting part of the proximal wing pouch from homozygous *ds^Cr+:V5^* (C), *ds*^*Cr*Δ*A:V5*^(F), and *ds*^*Cr*Δ*D:V5*^(I). **(J)** Western blot on lysates of third instar wing discs showing relative levels of D:V5 in the indicated genotypes. The two bands reflect Ds processing (Ambegaonkar et al., 2012). **(K)** Histogram showing quantification of western blots on Ds:V5 proteins, based on 9 replicates for *ds^Cr+:V5^*, *dsCr*^Δ*A:V5*^ and *dsCr*^Δ*D:V5*^, and 7 replicates for *ds^CrD^*^Δ*II:V5*^and *ds^CrD^*^Δ*III:V5*^. The statistical significance of differences as compared to *ds^Cr+:V5^* is indicated in green.

To compare Ds expression levels in different alleles, we conducted staining experiments in parallel and used identical imaging conditions. All of the *ds* alleles express Ds:V5 in a proximal-to-distal gradient (Figure 3D, G, Supplemental Figure S4). For *ds*^*Cr*Δ*B:V5*^; *ds*^*Cr*Δ*C:V5*^; *ds*^*Cr*Δ*E:V5*^ and *ds*^*Cr*Δ*F:V5*^, each of which provided essentially normal Ds activity, Ds:V5 protein appears to be expressed at the same levels and with the same localization as Ds:V5 expressed by *ds^Cr+:V5^* (Supplemental Figure S4A-L). For Ds:V5 expressed by *ds*^*Cr*Δ*A:V5*^, which provides enhanced Ds activity, protein levels appear increased relative to *ds^Cr+:V5^* (Figure 3D). In contrast, For Ds:V5 expressed from *ds*^*Cr*Δ*D:V5*^, which provides decreased Ds activity, protein levels appear reduced relative to *ds^Cr+:V5^* (Figure 3G). The increases and decreases in Ds:V5 staining were observed across different regions of the wing disc, including the wing pouch and the wing hinge (Figure 3D-I). Ds:V5 staining appears much reduced and poorly localized from *ds*^*Cr*Δ*ICD:V5*^ wing discs (Supplemental Figure S4M-O), which suggests that the ICD is important for normal localization and levels of Ds protein. Examination of Ds protein localization in wing imaginal discs from larvae with the *ds^CrDCHS1:V5^* hybrid transgene revealed that the Ds-DCHS1 hybrid protein is not properly localized. Rather than exhibiting normal accumulation at the subapical membrane, Ds:DCHS1:V5 was observed to accumulate inside cells, possibly due to mis-folding (Supplemental Figure S4P-R).

We also performed Western blot analysis on alleles for which immunostaining identified differences in expression. This confirmed that deleting the A motif leads to increased levels of Ds protein as compared to control *ds^Cr+:V5^* protein levels (Figure 3J, K). A slight decrease in Ds protein levels was detected by western blotting when the D motif was deleted, but it was not statistically significantly (Figure 3J, K). Thus, we infer that the reduced cell membrane staining detected in *ds*^*Cr*Δ*D:V5*^ wing discs primarily reflects mis-localization rather than decreased protein levels.

### The D motif of Ds is required for regulation by Lft

The decreased levels of Ds at cell membranes observed in *ds*^*Cr*Δ*D:V5*^ suggested that the D motif is required for regulation of Ds by a factor that interacts with it through this motif. The *lft* gene acts post-transcriptionally to promote normal membrane levels of both Fat and Ds in wing discs (Mao et al., 2009). To examine whether Lft regulation is mediated through the D motif, we compared Lft overexpression in control *ds^Cr+:V5^* wing discs and *ds*^*Cr*Δ*D:V5*^ wing discs. The UAS-Gal4 system was used to overexpress Lft in the posterior compartment, under control of a *hh-Gal4* driver. This enables comparison of Ds:V5 protein between posterior cells, which over-express Lft, and anterior cells, which have endogenous levels of Lft. In control *ds^Cr^*^+:V5^ discs, Lft over-expression increases visible levels of Ds:V5 as compared to anterior compartment staining (Figure 4A), consistent with prior studies (Mao et al., 2009). Conversely, in *ds*^*Cr*Δ*D:V5*^ discs, overexpression of Lft does not noticeably increase DsΔD:V5 protein as compared to levels in the anterior compartment (Figure 4B). This suggests that the reduced levels of Ds:V5 protein observed in *ds*^*Cr*Δ*D:V5*^ discs could be due to an inability to be positively regulated by Lft.

**Figure 4.**
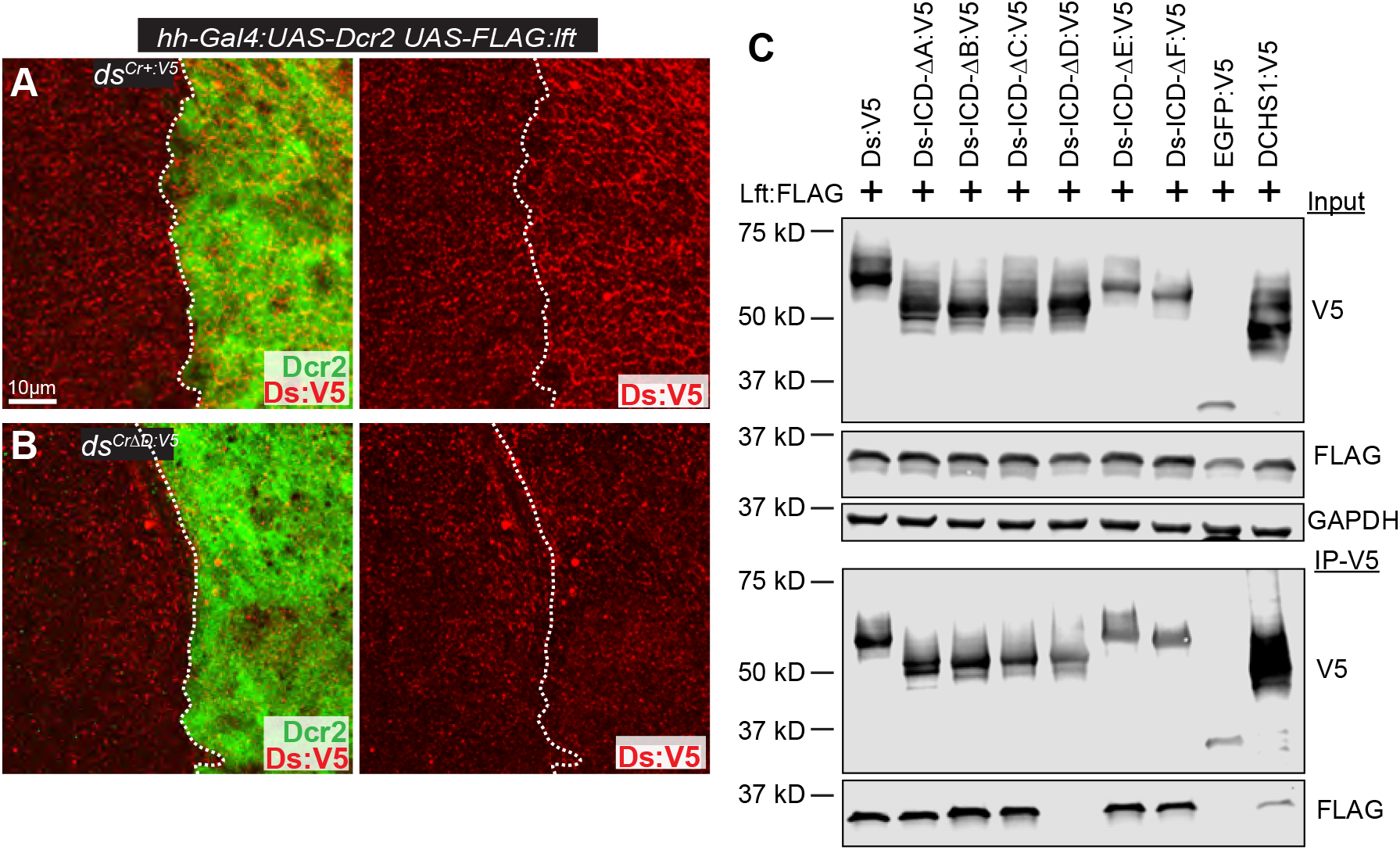
The D motif mediates regulation of Ds by Lft. **(A, B)** Wing discs expressing *hh*-*Gal4* driven *UAS*-*FLAG*:*lft* in (A) homozygous *ds^Cr+:V5^*, or (B) homozygous *ds*^*Cr*Δ*D:V5*^, with UAS-Dcr2 marking posterior (green), stained for V5 (red) to reveal Ds. **(C)** Western blot showing results of co-immunoprecipitation experiments between V5-tagged Ds and FLAG-tagged Lft expressed in S2 cells. Top three panels show blots of cell lysates transfected to express the indicated proteins, using antibodies indicated at right. GAPDH is a control for loading and transfer. Bottom two panels show blots on proteins immunoprecipitated with anti-V5 beads and detected with V5 or FLAG antibodies, as indicated.

To investigate whether this reflects direct association of Lft with Ds through this motif, we mapped regions of the ICD required for Lft binding. Earlier studies reported that Lft could associate with the Ds ICD but did not identify where within the ICD Lft binds (Mao et al., 2009). To address this, we expressed the ICDs of Ds motif deletion constructs in cultured *Drosophila* S2 cells, together with FLAG-tagged Lft. The intact Ds ICD could co-immunoprecipitate Lft. Deletion of motifs A, B, C, E, or F does not affect the ability of Lft to co-immunoprecipitate with the Ds-ICD. Conversely, deletion of the D motif (Ds-ICD-ΔD:V5) results in loss of Lft - Ds-ICD co-precipitation (Figure 4C). Thus, amongst conserved motifs in the Ds ICD, the D motif is uniquely required for association with Lft. Together with the insensitivity of the *ds*^*Cr*Δ*D:V5*^ mutant to overexpression of Lft, these results imply that the association of Lft with the D motif is required for Ds regulation.

The ability of Lft to associate with Fat and Ds is conserved by its mammalian homologues LIX1 and LIX1L (Mao et al., 2009), and we found that deletion of the D-motif also eliminates the binding of LIX1 and LIX1L to the Ds ICD (Supplemental Figure S5). Conservation across species of the ability of Ds and Lft proteins to bind to each is further supported by the observation that Lft is co-immunoprecipitated by the DCHS1 ICD (Figure 4C).

### The D motif of Ds is required for Dachs regulation and binding

The influence of Ds-Ft signaling on growth and polarity involves regulation of Dachs. Fat promotes removal of Dachs from cell membranes, driving the changes in levels of membrane Dachs that influence Hippo signaling and the polarization of Dachs that influences PCP (reviewed in Fulford and McNeill, 2020; Gridnev and Misra, 2022; Strutt and Strutt, 2021). Previous studies have also reported that Dachs can co-localize in puncta with Ds and physically associate with the Ds-ICD (Ambegaonkar et al., 2012; Bosveld et al., 2012; Brittle et al., 2012). However, the mechanistic significance of Ds-Dachs association to Dachs regulation remains unknown.

To investigate this, we generated mitotic clones for the Ds-ICD motif deletions that have significant phenotypes, and examined Dachs protein, using genomic GFP-labelled Dachs, in wing discs with these clones. Dachs is enriched at the subapical membrane in *ds*^*Cr*Δ*D:V5*^ mutant clones (Figure 5C), whereas Dachs levels appear unaffected in *ds^Cr^*^+*:V5*^ control clones (Figure 5A). Mitotic clones of *ds*^*CrΔA:V5*^ also do not have evident effects on Dachs levels or localization (Figure 5B). The increased accumulation of Dachs in *ds*^*Cr*Δ*D:V5*^ clones identifies the Ds-ICD D region as required for normal Dachs levels.

**Figure 5.**
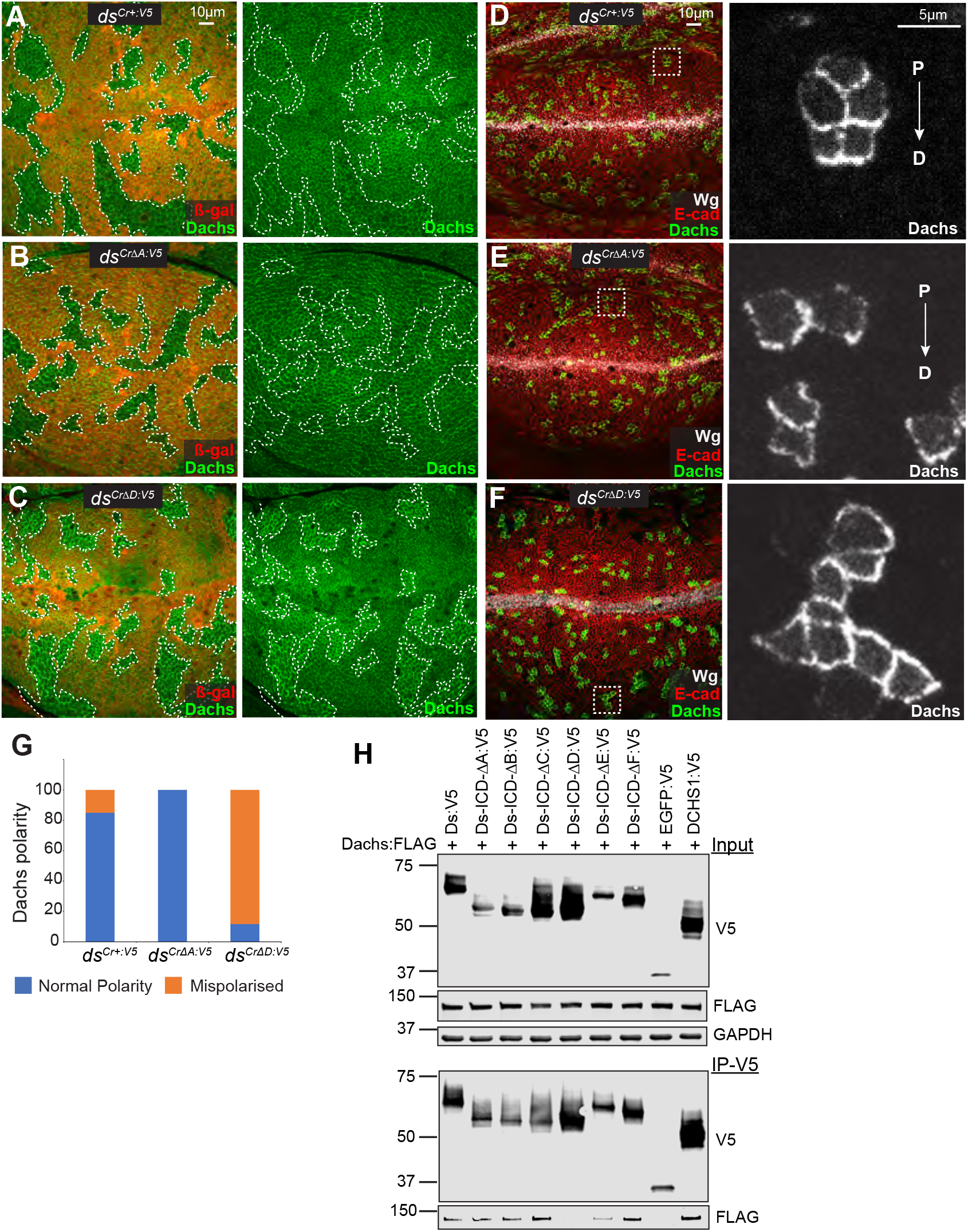
The Ds-D motif is required for Dachs regulation and binding. **(A-C)** Wing discs of *hs-Flp*; *ds^Cr+:V5^ FRT40A/arm-lacZ FRT40A; Dachs:GFP/+* **(A)**, *hs-Flp*; *d^sCr^*^Δ^*^A:V5^ FRT40A/arm-lacZ FRT40A; Dachs:GFP/+* **(B),** or *hs-Flp*; *ds^Cr^*^Δ^*^D:V5^ FRT40A/arm-lacZ FRT40A; Dachs:GFP/+* **(C)**, with mitotic clones homozygous for *ds^Cr+:V5^*(A), *d^sCr^*^Δ^*^A:V5^* (B), or *ds^Cr^*^Δ^*^D:V5^* (C), marked by loss of ß-gal (red), outlined with white dotted lines, to show effects on Dachs:GFP (green). **(D-F)** Wing discs of *hs-Flp*; *ds^Cr+:V5^/ds^ΔEx12^; Act>stop>EGFP:Dachs/+* **(D)***, hs-Flp*; *ds^Cr^*^Δ^*^A:V5^/ds^ΔEx12^; Act>stop>EGFP:Dachs/+* **(E)**, *or hs-Flp*; *ds^Cr^*^Δ^*^D:V5^/ds^ΔEx12^; Act>stop>EGFP:Dachs/+* **(F)** stained for Wg (white) and E-cad (red) and with Flip-out clones expressing Dachs:GFP (green/white). Panels at right show Dachs:GFP (white) in higher magnification images of the boxed regions. Proximal (P) to distal (D) orientation is indicated. **(G)** Histogram showing classification of Dachs polarity in Flp-out clones in different *ds* genotypes. 60 clones were scored per genotype. **(H)** Western blot showing results of co-immunoprecipitation experiments between V5-tagged Ds proteins and FLAG-tagged Dachs transfected into S2 cells. Top three panels show blots on total cell lysates transfected to express the indicated proteins, using the antibodies indicated at right. Bottom two panels show blots on proteins immunoprecipitated with anti-V5 beads and detected with V5 or FLAG antibodies.

To determine the influence of Ds-ICD motif deletions on Dachs polarization, we made small Flp-out clones expressing GFP-tagged Dachs, which facilitates visualization of polarization. A Flip-out cassette expressing Dachs:GFP under the control of the Actin-5C promoter, with an intervening transcriptional stop cassette flanked by FRT sites, was introduced into Ds-ICD motif deletion backgrounds. A pulse of heat-shock-induced Flipase expression led to Dachs:GFP-expressing clones. Most clones in *ds^Cr^*^+*:V5*^ wing discs exhibit normal, distally-oriented Dachs polarity (Figure 5D, G). Dachs is also normally polarized in *ds*^*CrΔA:V5*^ wing discs (Figure 5E, G). Conversely, most Dachs:GFP clones in *ds*^*Cr*Δ*D:V5*^ mutant wing discs have mispolarised Dachs (Figure 5F, G). This mispolarization is consistent with the observation that PCP is abnormal in *ds*^*Cr*Δ*D:V5*^ mutant wing discs.

We also mapped regions of the Ds ICD required for association with Dachs through co-immunoprecipitation experiments, using a Flag-tagged Dachs construct and V5-tagged Ds ICD constructs. Deletion of the A, B, C, E, or F motifs did not impair the ability to bind Dachs. Conversely, deletion of the D-motif abolished detectable Dachs binding (Figure 5H). We also found that V5-tagged DCHS1 can co-precipitate Dachs, consistent with the conclusion that Ds-Dachs association is mediated by a conserved motif (Figure 5H).

### The D motif of Ds is required for Sple binding

The Sple isoform of the Pk-Sple locus links Ds-Fat PCP with Fz-PCP and has been reported to physically associate with Ds and Dachs, and to co-localize in puncta with Ds and Dachs (Ambegaonkar and Irvine, 2015; Ayukawa et al., 2014; Gubb et al., 1999; Merkel et al., 2014; Olofsson et al., 2014). To investigate the role of the Ds-ICD in Sple regulation, we determined whether any conserved motifs are required for association with Sple by performing co-immunoprecipitation experiments on lysates of S2 cells expressing co-transfected FLAG-tagged Sple-N terminal region and V5-tagged Ds-ICD constructs. Deletions of the A, B, C, E, or F motifs did not impair the ability of the Ds ICD to bind Sple-N. Conversely, deletion of the D motif eliminated detectable binding to Sple-N (Supplemental Figure S6), implying that Sple associates with Ds through the D-motif of the ICD. This conclusion is potentially consistent with the observation that *ds*^*Cr*Δ*D:V5*^ is the only one of the motif deletions that exhibits significant PCP phenotypes (Figure 2C, E). However, as motif D also influences levels of Ds and localization of Dachs, the PCP phenotypes observed do not necessarily stem directly from impairment of association with Sple.

### The D motif of Ds is sufficient for association with Lft, Dachs, and Sple

The observation that each of three different binding partners of Ds: Lft, Dachs, and Sple, all require the D motif for association with Ds raised concerns about the specificity of this requirement. For example, if deletion of the D motif results in misfolded protein, then the requirement might be indirect. As an approach to address this, we investigated whether the D motif is sufficient, for interaction with Lft, Dachs, and Sple. As the D motif is only 64 amino acids, this was achieved by creating and expressing a Ds ICD D-motif:EGFP:V5 fusion protein, which was then used in co-immunoprecipitation experiments, alongside EGFP:V5 as a negative control. This revealed that Ds-D+EGFP:V5 could specifically co-immunoprecipitate Lft in S2 cells, and thus that the D motif is sufficient for Lft binding (Figure 6A). Similarly, Ds-D+EGFP:V5, but not EGFP:V5 could co-immunoprecipitate Dachs and Sple (Figure 6B, C). Thus, the D motif is not only necessary, but also sufficient, for association of Ds with Lft, Dachs, and Sple.

**Figure 6.**
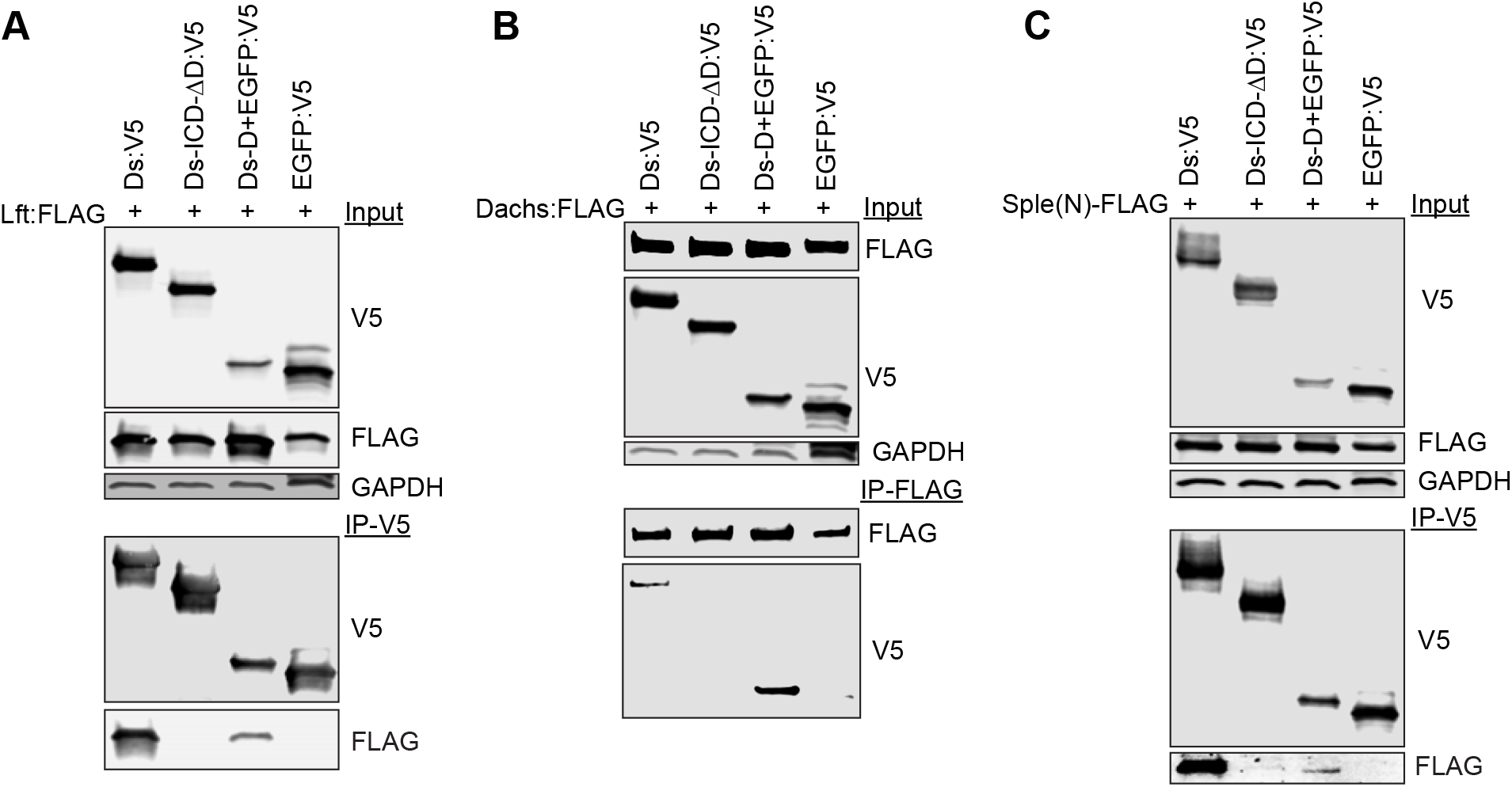
The Ds-D region is sufficient for Lft, Dachs and Sple-N association. Western blot showing results of the co-immunoprecipitation experiments between V5-tagged Ds and FLAG-tagged Lft (A), Dachs (B), or Sple-N (C). Top three panels show blots on cell lysates transfected to express the indicated proteins, using antibodies indicated at right. Bottom two panels show blots on proteins immunoprecipitated with anti-V5 beads and detected with V5 or FLAG antibodies.

### Partial separation of Lft and Dachs binding within the Ds ICD

We next investigated whether any of the distinct functions of the D motif could be separated. Structural prediction using AlphaFold (Jumper et al., 2021) suggests that around 76% of the ICD is intrinsically disordered, but structured regions including predicted alpha helices occur in motifs B, C, and D (Figure 7A). If Lft, Dachs, and Sple have differential binding regions within the D-motif, this might be revealed by smaller deletions. Using the predicted structure as a guide, we thus subdivided the D motif into three smaller regions, which we refer to as D-I, D-II, and D-III (Figure 7A, B). We note that a region of the Fat-ICD with sequence similarity to the Ds-ICD overlaps the D motif (Clark et al., 1995; Mao et al., 2009), with maximum similarity in the D-II and D-III regions (Figure 7C).

**Figure 7.**
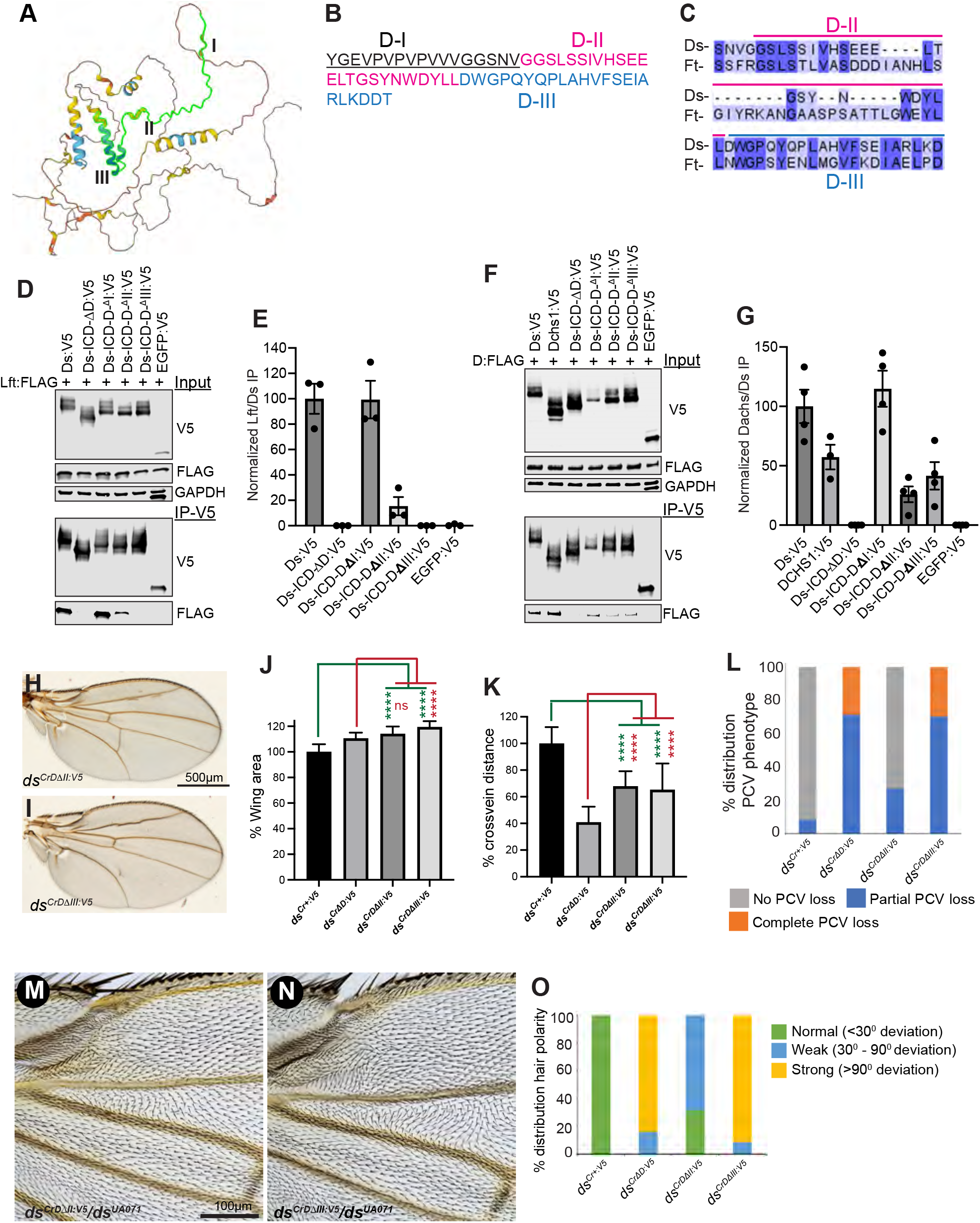
Separating functions encoded by the Ds D region. **(A)** Predicted structure of the Ds-ICD. The D motif is highlighted in green, with regions I, II and III indicated. **(B)** Amino acid sequence of the D motif, with regions I, II, and III in black, red, and blue letters, respectively. **(C)** Sequence comparison between a portion of the Fat-ICD and the Ds-D region (Clark et al., 1995; Mao et al., 2009). **(D,F)** Western blots showing results of co-immunoprecipitation experiments between V5-tagged Ds proteins and FLAG-tagged Lft **(D)**, or FLAG-tagged Dachs **(E)**, co-expressed in S2 cells. Top three panels show blots of cell lysates transfected to express the indicated proteins, detected using antibodies indicated at right. Bottom two panels show blots on proteins immunoprecipitated with anti-V5 beads and detected with V5 and FLAG antibody, as indicated. **(E,G)** Quantitation of relative co-precipitation of Lft (E, based on 3 replicates) or Dachs (G, based on 4 replicates) with the indicated Ds-ICD constructs. **(H, I)** Adult male wings from flies with Ds-ICD D-II deletion (*ds^CrD^*^Δ*II:V5*^*/ds^UA071^*) (H), or D-III deletion (*ds^CrD^*^Δ*III:V5*^*/ds^UA071^*) (I). **(J-L, O)** Histograms representing relative wing area (J), relative cross-vein distance (K), distribution of PCV loss (L) and distribution of wing PCP phenotypes (O) from the given *ds* alleles over *ds^UA071^*. Each bar indicates mean ± SD from measurement of n=24 (*ds^CrD^*^Δ*II:V5*^) or n=25 wings (*ds^CrD^*^Δ*III:V5*^). **(M, N)** Proximal anterior region of adult male wings from *ds^CrD^*^Δ*II:V5*^*/ds^UA071^* (M), and *ds^CrD^*^Δ*III:V5*^*/ds^UA071^* (N). Scale bar = 100 µm.

Constructs with smaller deletions within the D-motif (D-I, D-II, D-III) were expressed in S2 cells together with tagged Lft, Dachs, or Sple. Co-immunoprecipitation experiments revealed that removing the D-I region does not affect binding of the Ds-ICD to Lft (Figure 7D,E). However, deleting the D-II region reduced binding of Lft to 15% of Ds-ICD full-length levels (Figure 7D,E), and deleting the D-III region completely eliminated binding of Lft to the Ds-ICD (Figure 7D,E). Association of Dachs with the Ds-ICD was unaffected by deletion of the D-I region, but reduced to 26% of Ds-ICD full-length levels by deletion of the D-II region, and to 42% by deletion of the D-III region (Figure 7F,G). Association of Sple with the Ds-ICD was reduced to around 20% of Ds-ICD full-length levels by deletion of D-II or D-III (Supplemental Figure S6C,D). Altogether then, we could partially separate the sequence motifs required for association of Lft and Dachs with Ds, as the D-II region is preferentially required for association with Dachs and the D-III region is absolutely required for association with Lft.

Based these co-immunoprecipitation experiments, we then used RMCE to generate flies with genomic deletions of the D-II or D-III regions, and compared their phenotypes to those of *ds^Cr^*^+*:V5*^ and *ds*^*Cr*Δ*D:V5*^. Deletion of the D-II region, *ds^CrD^*^*ΔII:V5*^, which reduces Dachs, Sple and Lft binding, leads to an increase in wing size similar to that *ds*^*Cr*Δ*D:V5*^ flies (Figure 7H, J). Relative cross-vein distance was decreased compared to *ds^Cr^*^+*:V5*^, but less so than observed in *ds*^*Cr*Δ*D:V5*^ flies (Figure 7K). Loss of the PCV, and disruption of wing hair polarity were observed, but these phenotypes were substantially weaker than the phenotypes in *ds*^*Cr*Δ*D:V5*^ flies (Figure 7L-O). These observations suggest that some functions of the Ds ICD, related to its contributions to wing growth, are severely compromised by the D-II deletion, but other functions, including those related to PCP, are only partially compromised.

Deletion of the D-III region, *ds^CrD^*^*ΔIII:V5*^, which eliminates Lft binding and reduces Dachs binding, leads to an increase in wing size that is even greater than that observed in *ds*^*Cr*Δ*D:V5*^ flies (19% versus 11% larger than controls, (Figure 7I, J). Loss of the PCV and disruption of wing hair polarity phenotypes were also observed and appear similar in strength to the complete deletion of the D motif (Figure 7L-O). D-III deletion wings also show a decrease in relative cross-vein distance (Figure 7I, K), although the phenotype was milder than observed with the full D motif deletion. Overall, removal of the D-III region results in phenotypes that are similar to those observed in *ds*^*Cr*Δ*D:V5*^ flies.

To investigate the molecular consequences of these smaller deletions, we examined Ds protein levels and localization through immunostaining and western blotting. This revealed that Ds protein localization in D-II deletion mutants appears overall similar to that in *ds^Cr^*^+*:V5*^ control flies (Figure 8A-C). Conversely, Ds protein localization in D-III deletion mutants showed reduced Ds localization to cell junctions, similar to that observed in *ds*^*Cr*Δ*D:V5*^ flies (Figure 8D-F). As for *ds*^*Cr*Δ*D:V5*^, deleting either the D-II or D-III motifs did not significantly alter Ds protein levels as assayed by western blotting (Figure 3 J,K).

**Figure 8.**
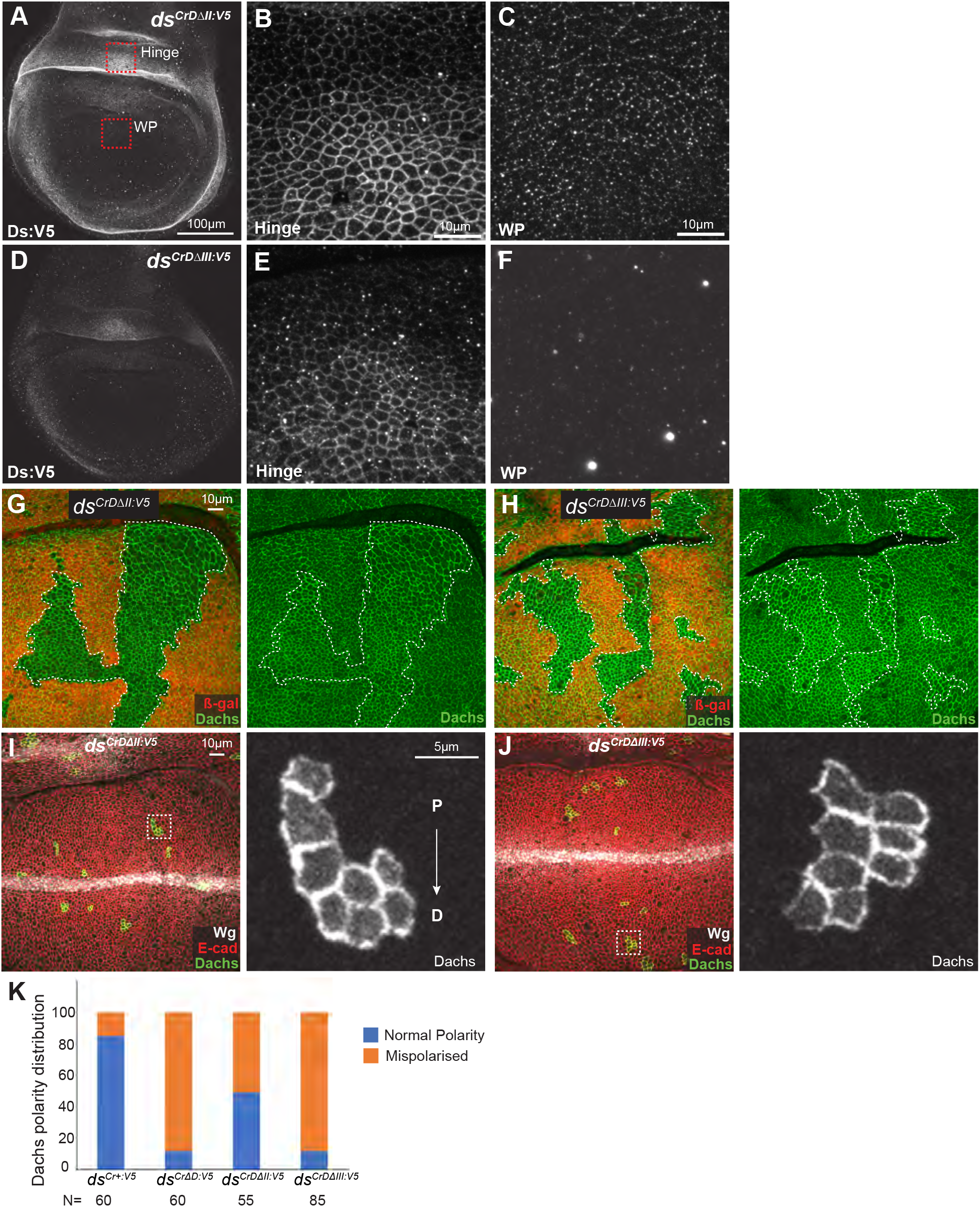
Influence of D-II and D-III Ds-ICD deletions on Ds and Dachs. **(A-F)** Wing discs from homozygous *ds^CrD^*^Δ^*^II:V5^* (A) or *ds^CrD^*^Δ^*^III:V5^* (D) stained for Ds:V5. Red square boxes in (A) show approximate locations in the hinge and wing pouch (WP) corresponding to the higher magnification panels at right. (**A,D**) Lower magnification images showing the wing pouch and hinge regions, Scale bar = 100 µm. **(B,C,E,F)** Higher magnification images depicting part of the wing hinge (**B,E**) or proximal wing pouch (**C,F**) in discs expressing V5-tagged Ds from *ds^CrD^*^Δ^*^II:V5^* (**B,C**) or *ds^CrD^*^Δ^*^III:V5^* (**E,F**). Scale bar = 10 µm. **(G,H)** Wing discs from *hs-Flp*; *ds^CrD^*^Δ^*^II:V5^ FRT40A/arm-lacZ FRT40A; Dachs:GFP/+* **(G)**, or *hs-Flp*; *ds^CrD^*^Δ^*^III:V5^ FRT40A/arm-lacZ FRT40A; Dachs:GFP/+* **(H),** with mitotic clones homozygous for *ds^CrD^*^Δ^*^II:V5^* (G), or *ds^CrD^*^Δ^*^III:V5^* (H), marked by loss of ß-gal (red), outlined with white dotted lines, to show effects on Dachs:GFP (green). **(I,J)** Wing discs from *hs-Flp*; *ds^CrD^*^Δ^*^II:V5^/ds^ΔEx12^; Act>stop>EGFP:Dachs/+* **(I)**, or *hs-Flp*; *ds^CrD^*^Δ^*^III:V5^/ds^ΔEx12^; Act>stop>EGFP:Dachs/+* **(J)**, stained for expression of Wg (white) and E-cad (red) and with Flip-out clones expressing Dachs:GFP (green/white). Panels at right show Dachs:GFP (white) in higher magnification images of the boxed regions. Proximal (P) to distal (D) orientation is indicated. **(K)** Histogram showing the classification of Dachs polarity in Flp-out clones in different *ds* genotypes as normal (blue color) or mispolarised (orange). N= number scored.

We next examined the effects of the D-II and D-III deletions on Dachs levels and localization. Mitotic clones homozygous for *ds*^*Cr*Δ*D-II:V5*^ or *ds*^*Cr*Δ*D-III:V5*^ in developing wing discs revealed that Dachs is elevated at the subapical membrane by both D-II and D-III motif deletions as compared Dachs in neighboring cells (Figure 8G, H). The elevated levels of Dachs are consistent with the observation that both of these mutations are associated with increased wing growth, as membrane levels of Dachs correlate with its downregulation of Wts in the Fat-Hippo pathway (Brittle et al., 2012; Mao et al., 2006; Rogulja et al., 2008; Vrabioiu and Struhl, 2015). Flp-out clones expressing Dachs:GFP were examined to assess Dachs polarization. This revealed that Dachs polarity was often normal in flies with the D-II deletion (Figure 8I, K), but Dachs was mostly mis-polarized in flies with the D-III deletion (Figure 8J, K). The distinct effects of these different small deletions on Dachs polarity correlates with their distinct effects on wing hair PCP, CV spacing, and PCV loss, which are strongly affected by the D-III deletion but only mildly affected by the D-II deletion.

### Influence of Ds ICD deletions on Hindgut looping and MyoID binding

In addition to its Dachs-dependent roles in Hippo signaling and PCP, Ds is also required for normal looping of the hindgut (González-Morales et al., 2015). In wild-type flies, the adult hindgut exhibits a clockwise coiling pattern (dextral looping), forming a single stereotyped loop that is located on the right side of the abdomen when viewed dorsally. Flies mutant for the myosin family protein MyoID exhibit a L-R patterning defect, in which hindgut looping is predominantly sinistral (Hozumi et al., 2006). *ds* mutant flies have a mis-looped phenotype (Figure 9B, I), which is thought to be due to mis-regulation of MyoID, which the Ds-ICD can physically associate with (González-Morales et al., 2015).

**Figure 9.**
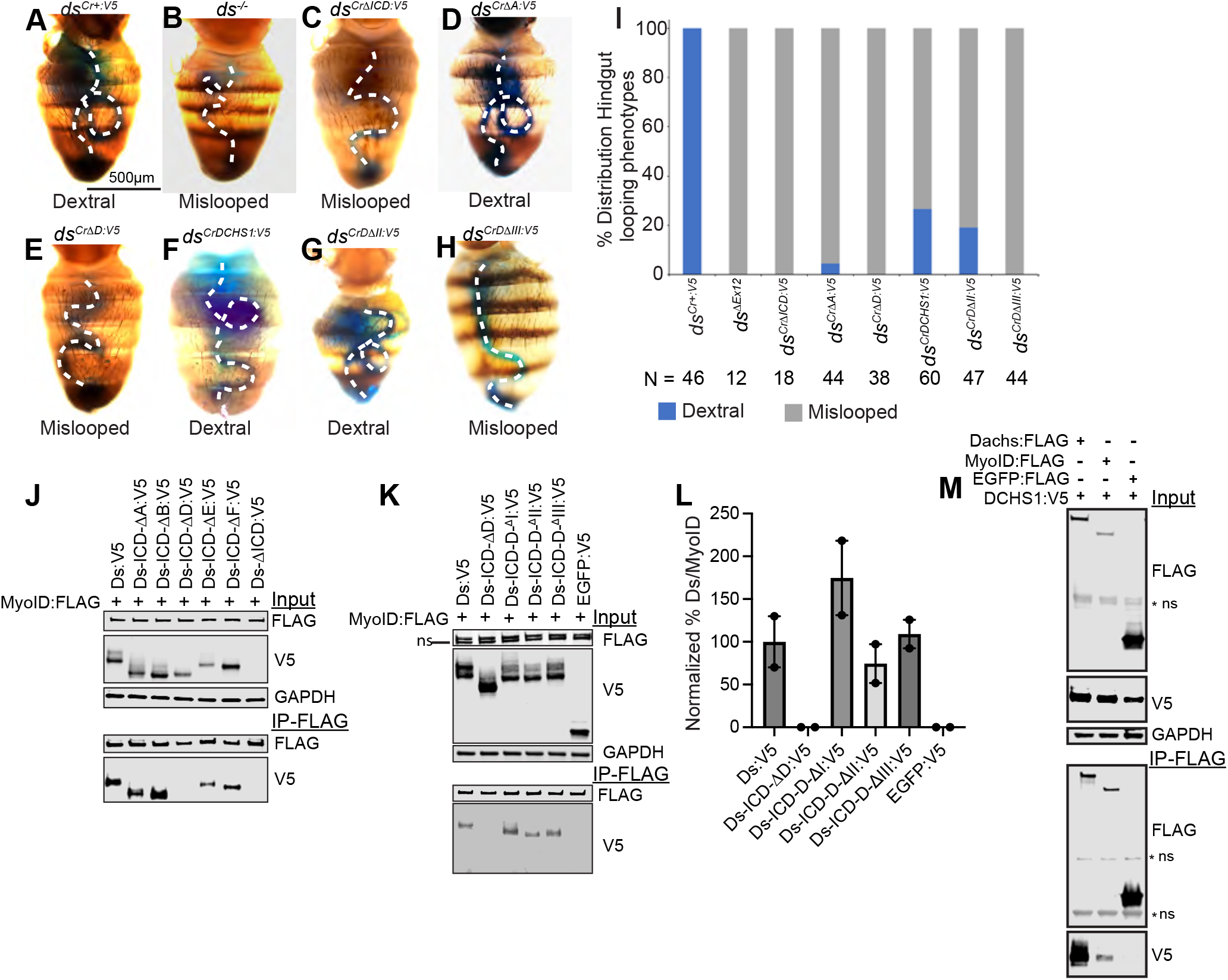
The D motif is required for dextral hindgut looping and MyoID binding. **(A-H)** Dorsal views of adult abdomens fed blue dye to mark the hindgut. (A) *ds^Cr+:V5^/ds^UA071^* (control), (B) *ds^36D^/ds^UA071^,* (C) *ds*^*Cr*Δ*ICD:V5*^*/ds^UA071^*, (D) *ds*^*Cr*Δ*A:V5*^*/ds^UA071^,* (E) *ds*^*Cr*Δ*D:V5*^*/ds^UA071^,* (F) *ds^CrDCHS1:V5^/ds^UA071^,* (G) *ds^CrD^*Δ*II:V5 /ds^UA071^,* (H) *ds^CrD^*Δ*III:V5 /ds^UA071^.* (I) Histogram showing distribution of phenotypes among different genotypes, scored as normal dextral looping (blue), or mislooped (gray). N = number of flies scored. **(J,K)** Western blots showing results of co-immunoprecipitation experiments between V5-tagged Ds proteins and FLAG-tagged MyoID. Top three panels show blots on total cell lysates transfected to express the indicated proteins, using the antibodies indicated at right. Bottom two panels show blots on proteins immunoprecipitated with anti-V5 beads and detected with V5 or FLAG antibodies. The lower band on the input FLAG blot is non-specific (ns). **(L)** Quantitation of relative co-precipitation of MyoID with the indicated Ds-ICD:V5 constructs. **(M)** Western blot showing results of co-immunoprecipitation experiments between V5-tagged DCHS1 ICD and the indicated FLAG-tagged proteins. Top three panels show blots on total cell lysates transfected to express the indicated proteins, using the antibodies indicated at right. Bottom two panels show blots on proteins immunoprecipitated with anti-FLAG beads and detected with V5 or FLAG antibodies. Some non-specific (ns) bands are indicated.

Looping of the hindgut can be visualized using ingestion of food with blue dye, which labels the gut lumen. We used this to examine hindgut looping in several Ds-ICD deletion mutants. Control *ds^Cr^*^+*:V5*^ flies all exhibit normal dextral looping (Figure 9A, I). Deletion of the A-motif does not result in significant hindgut looping defects (Figure 9D, I). Conversely, complete ICD deletion, or deletion of just the D-motif, results in a completely penetrant mislooped phenotype like that of *ds* mutants (Figure 9C, E, I). The smaller D-III deletion mutant also exhibit a completely penetrant mis-looped phenotype (Figure 9H, I), but the D-II deletion exhibits a milder phenotype, with 19% of flies exhibiting normal dextral looping (Figure 9G, I). The *ds^Cr^*^DCHS1*:V5*^ hybrid transgene also exhibited partial Ds activity in this assay, with normal dextral looping observed in 27% of flies and mis-looping in the remaining 73% (Figure 9F, I).

MyoID was reported to bind to the Ds-ICD, but specific regions responsible for this binding were not identified (González-Morales et al., 2015). Thus, we performed co-immunoprecipitation experiments on lysates of S2 cells transfected to express Flag-tagged MyoID and V5-tagged Ds-ICD constructs (Figure 9J). This established that deletion of the D-motif specifically eliminated

MyoID binding, which is consistent with observations that the D motif disrupts normal hindgut looping. Using smaller deletions within the D region, we found that the D-II deletion reduced, but does not eliminate, binding to Ds, whereas the D-I and D-III deletions did not reduce binding (Figure 9K,L). The ability to associate with MyoID was conserved in the DCHS1 ICD (Fig. 9M). Like Dachs, MyoID is an atypical myosin protein, so their association with the same region of the ICD suggests that this association is mediated through aspects of myosin structure that are common to both proteins.

## Discussion

Dissection of the Fat ICD provided insight into how it mediates downstream signal transduction (Matakatsu and Blair, 2012; Pan et al., 2013; Zhao et al., 2013). In the case of Fat, multiple distinct motifs carry out different aspects of its activity. Here, we initiated a comparable study of the Ds ICD, examining the functional requirements for different sequence motifs, and assessing the ability of some known Ds interactors to associate with these motifs, in order to define roles of the Ds ICD and investigate the hypothesis that Ds mediates downstream signal transduction. Amino acid sequence comparisons identified several conserved regions, which we divided into six motifs, four conserved to vertebrates and two only conserved within insects. Evolutionary conservation between insects and vertebrates implies that the motifs are functionally essential, but multiple phenotypic assays identified only one of the four motifs conserved in vertebrates, the D motif, as having major effects on Ds activity, whereas deletion of the B, E, or F motifs had either no or only minor effects on Ds activity. We also attempted to address the significance of evolutionary conservation of the Ds ICD by assaying the ability of the human DCHS1 ICD to provide Ds ICD function in vivo, using a hybrid transgene encoding the ECD and TM of *Drosophila* Ds and the ICD of human DCHS1. However, imaging revealed that the hybrid protein is mis-localized, suggesting that it may be mostly misfolded, which could explain its failure to provide significant Ds activity in vivo. Nonetheless, co-immunoprecipitation experiments confirmed that key partners of *Drosophila* Ds, including Lft, Dachs, MyoID and Sple, could associate with the DCHS1 ICD in cultured cells.

The two motifs that had substantial effects on Ds activity, the A and D motifs, each affected Ds protein levels or membrane localization, and consequently a significant part of the effect of these motifs on Ds function could be due to their effects on the levels of Ds at its normal location. Thus, we cannot attribute specific roles for these sequence motifs in downstream signaling, and indeed their effects might be primarily due to increased or decreased activation of Fat by Ds. Importantly then, deletion of the smaller D-II motif impairs Ds function without noticeably decreasing Ds levels. The lack of other identifiable deletions impairing Ds function is somewhat surprising if Ds functions as a signal transducing receptor, but it could be that some functions are provided by larger regions in a redundant fashion, such that larger deletions would be needed to impair function. Alternatively, it could be that Ds functions primarily as ligand, with the principal function of the ICD being to modulate the levels and localization of the ECD so that it can appropriately regulate Fat. We also note that as motifs that our molecular studies indicate should be crucial for PCP, ie interaction with Dachs and Sple, are embedded within a region that is also crucial for regulation of Ds levels, we could not fully separate potential PCP activities downstream of Ds from regulation of PCP via Fat.

Our studies have provided a molecular explanation for part of the impact of the D motif deletion on Ds by the discovery that this motif is necessary and sufficient for mediating physical association with Lft. Moreover, deletion of the D motif renders Ds insensitive to increased Lft expression. Lft was previously identified as being required for normal levels and distribution of Fat and Ds (Mao et al., 2009). Intriguingly, there is a region of sequence similarity between the Fat and Ds ICDs that partially overlaps with the Ds D region, which likely corresponds to a shared Lft-interacting sequence motif. This motif also has some similarity to sequences within the E-cadherin ICD (Clark et al., 1995; Mao et al., 2009). We emphasize though, that the phenotype of animals with deletion of the Ds D region, or even of the smaller D-III region, is more severe than the phenotype of *lft* mutants. This implies that the D and D-III regions have other functions beyond mediating association with Lft.

One such function that we identified is interaction with Dachs, as co-immunoprecipitation experiments established that the D region is necessary and sufficient for association of the Ds ICD with Dachs. However, our studies also suggest that the ability of Ds to associate with Dachs is not required for Dachs membrane localization or polarization, both of which are preserved in D-II deletion mutants, despite a significant reduction (74%) in Dachs binding. This supports the hypothesis that Dachs regulation is primarily mediated through the activity of the Fat ICD in antagonizing Dachs membrane localization rather than any ability of the Ds ICD to promote Dachs membrane localization. A region of zebrafish Dchs1b that interacts with the microtubule regulator Ttc28A overlaps with parts of the D-II and D-III regions (Chen et al 2018), so this could potentially be another factor affected by D region deletions.

The D motif is also required for association of Ds with MyoID, which suggests that this region could function as a common motif for interaction of unconventional myosins with Ds. This is intriguing as Dachs is not conserved in mammals, and we lack a good understanding of how Dchs1-Fat4 signaling regulates PCP in mammals. MyoID has been linked to L-R patterning in both vertebrates and invertebrates (Hozumi et al., 2006; Juan et al., 2018; Saydmohammed et al., 2018), and the observation that both Dachs and MyoID interact with the same, conserved, region of the Ds ICD supports the idea that myosin family proteins interacting with this motif could contribute to Dchs1-Fat4 modulation of PCP.

The mechanism by which the D-II region influences Ds function is not clear. The levels and localization of Ds appear similar to wild-type protein, PCP is only mildly affected, but the increased wing size and elevated membrane Dachs levels associated with this mutation indicate that Ds activity is compromised. As it is hard to understand how reduced Ds-Dachs binding would lead directly to elevated Dachs levels, we favor the hypothesis that this allele is defective in activating Fat. Molecular mechanisms associated with Fat activation by Ds have not been well defined. That is, activation of Fat involves binding by Ds, but whether Ds activates Fat by triggering Fat multimerization, conformational changes in Fat, recruitment of other proteins, or through other mechanisms remains uncertain. The *ds*^*Cr*Δ*D-II:V5*^ allele may thus prove useful for future investigations of this question.

## MATERIALS AND METHODS

### Drosophila genetics

Fly crosses were performed at 25°C unless otherwise noted. The following previously described fly stocks were used: *w^1118^, ds^UA071^* (Adler et al., 1998), *ds^36D^* (Rodriguez, 2004), *Df(2L)ED87/SM6a* (Bloomington #8677)*, UAS-FLAG:Lft* (Mao et al., 2009), *hh-GAL4 (FBti0017278), Dachs:GFP* (Bosveld et al., 2012)*, arm-lacZ* (Bloomington #7116), *Act>stop>EGFP:Dachs* (Brittle et al., 2012). *ds*^*ΔEx12*^*, ds^Cr+:V5^*, *ds^Cr+^*, *ds*^*CrΔA:V5*^*, ds^Cr^*^*ΔB:V5*^*, ds^Cr^*^*ΔC:V5*^, *ds*^*CrΔD:V5*^, *ds*^*CrDΔII:V5*^, *ds*^*CrDΔIII:V5*^, *ds*^*CrΔE:V5*^, *ds*^*CrΔF:V5*^, *ds*^*CrΔICD:V5*^, and *ds*^*CrDCHS1:V5*^ were generated in this study. The *ds*^Δ*Ex12*^ allele was generated by CRISPR/Cas9-mediated genome editing and homology-dependent repair, using services of WellGenetics. Two guide RNAs (CRISPR Target Site 01: AGTTCAGTAGCTGAAAAGGATGG and CRISPR Target Site 02: ACACGGATGTAATCGAGCACTGG) were used to delete *ds* Exon 12 and replaced with an RMCE cassette containing an attP cassette and a 3xP3-RFP selection marker. The RMCE cassette within *ds* was subsequently substituted via site-specific recombination with either wild-type *ds* sequences or with *ds* sequences in which one of the conserved motifs was deleted, or the entire *ds* ICD was deleted. We also generated additional fly lines in which the entire Ds-ICD was substituted with the ICD from human DCHS1 using microinjection services from BestGene. *ds* alleles were recombined with FRT40A for mitotic recombination. Mutant clones were generated at 72-84 hours AEL using *hs-FLP*, induced for 1 hour at 37°C, and analyzed 48 hours later. To make FLP-out Dachs:GFP clones, animals were cultured at 18*°C* for 7 days, heat-shocked 5 minutes at 35°C, and then wing discs were dissected out 18-24 hours later.

### Analysis of adult phenotypes

Adult male wings were dissected and mounted in Gary’s Magic Mount (4:1 Canada Balsam:Methyl Salicylate). Wings from at least 15 flies were photographed using a Zeiss Axioplan2 microscope and a Progress camera. Digital images of wings were traced manually to measure area and roundness using Fiji (Schindelin et al., 2012). Values were normalized to the average in controls and plotted by using GraphPad Prism. Cross-vein spacing was calculated by tracing the length of vein L4 between cross-veins, divided by the length of vein L3. For wings with incomplete cross-veins, we approximated the crossing points on the L4 vein using the incomplete cross-vein. Hair polarity was scored in the anterior proximal region of the wing based on the angle of deviation from the normal axis and categorized as normal (<30°), weak (30°-90°), or strong (>90°) if 10% or more wing hairs showed a deviation. PCV loss was scored manually and categorized as follows: No phenotype, incomplete PCV loss, and complete PCV loss. To examine hindgut looping, newly eclosed flies (both male and female) were fed fly food containing 2.5% Erioglaucine (Sigma, #861146), and the hindgut was examined through the cuticle (Gonzalez-Morales et al., 2015).

### Immunostaining wing imaginal discs

Wing discs were dissected from third instar larve at 96–120 h AEL, fixed for 15 minutes in 4% PFA and washed and stained as described previously (Rauskolb and Irvine, 2019). Images were collected on a Leica SP8 confocal microscope. For comparison of expression levels amongst different genotypes, consistent magnification, resolution, laser power, and detector settings were used. Primary antibodies were used for staining were: Mouse anti-V5 (Invitrogen R960-25; preabsorbed at 1:10 dilution, then used at 1:50), Rat anti-E-cadherin (DSHB DCAD2-c 1:200), Chicken anti-ß-gal (Abcam ab9361; 1:100), Rat anti-Fat (1:1500) (Feng and Irvine, 2009), Mouse anti-Wg (DSHB 9A4; 1:300), Rabbit anti-Dcr2 (Abcam ab4732, 1:1600), Rabbit anti-FLAG (Sigma F3165; 1:200). Secondary antibodies were from Jackson ImmunoResearch Laboratories, Invitrogen and Biotium (#20137). DNA was labelled with Hoechst 33342 (Life Technologies).

### Sequence analysis

Clustal Omega (https://www.ebi.ac.uk/jdispatcher/msa/clustalo) was used to identify sequence conservation among Dachsous ICDs from different species (Madeira et al., 2022), which was then displayed using Jalview (https://www.jalview.org) (Waterhouse et al., 2009). Dachsous-ICD structure was predicted by AlphaFold (https://alphafold.ebi.ac.uk) (Jumper et al., 2021).

### Cell culture, Immunoprecipitation and Western blotting

S2 cells were cultured in Schneider’s *Drosophila* Medium (Gibco # 21720001), supplemented with 10% FBS (ATCC #30-2020) and antibiotics (Gibco # 15240062), at 28°C. Cells were transfected using Effectene (Qiagen #301427), incubated for 44–48 hours at 28°C. Plasmids used to transfect S2 cells included: Aw-GAL4, pUAST-FLAG:Lft, pUAST-FLAG:LIX1, pUAST-FLAG:LIX1L (Mao et al., 2009), pUAST-Dachs-:FLAG (Mao et al., 2006), pUAST-GFP:V5, pUAST-Ds-TM-ICD:V5 (Mao et al., 2009), pUAST-Sple-N:FLAG (Ambegaonkar and Irvine, 2015). FLAG-tagged MyoID was generated for this study by cloning cDNA clone SD01662 from Drosophila Genomics Research Center (DGRC) into a pUAST vector using NEBuilder HiFi DNA Assembly Master Mix (NEB # E2621L). Plasmids encoding different Ds-ICD motif deletion constructs with a C-terminal V5-tag were created in this study: pUAST-Ds-TM-ICD-ΔA:V5, pUAST-Ds-TM-ICD-ΔB:V5, pUAST-Ds-TM-ICD-ΔC:V5, pUAST-Ds-TM-ICD-ΔD:V5, pUAST-Ds-TM-ICD-ΔE:V5, pUAST-Ds-TM-ICD-ΔF:V5, pUAST-Ds-TM-ICD-ΔD-II:V5, pUAST-Ds-TM-ICD-ΔD-III:V5, pUAST-Ds-TM-DCHS1-ICD:V5 and pUAST-Ds-D+EGFP:V5. pUAST-Ds-TM-ICD:V5 (Mao et al., 2009) was used to PCR amplify the Fat signal peptide along with the part of the Ds-TM region by using the following primer set: Fwd Primer: GAACTCTGAATAGGGAATTGGGATGGAGAGGCTACTGCTCC, Rev Primer: ATGGATGAATAGAAAGATTCC. Plasmids containing ds-Exon12 with different motifs deletion were provided by Wellgenetics. These plasmids were used as a template to amplify Ds-ICD sequence with different conserved motif deletions by using the following primer set: Fwd Primer: GGCAATTGGTCTACTGGTAGC, Rev Primer: GGAAATGTGGGGACACGGATGGGCGGAGGCGGATCCGGAAAACCCATCCCAAACCCCCTCTTGGGTTTGGACAGCACTCGTACCGGTCATCATCACCATCACCATTAAAATTCGTTAACAGATCTGCG. The above two sets of PCR products were gel purified and cloned in a pUAST-attB vector digested with EcoRI using NEBuilder HiFi DNA Assembly Master Mix (NEB # E2621L). Transfected cells were lysed in RIPA buffer (140mM NaCl, 10mM Tris-HCl pH 8.0, 1mM EDTA pH 8.0, 1.0% Triton X-100, 0.1% SDS, and 0.1% Sodium deoxycholate) supplemented with Protease Inhibitor Cocktail COMPLETE EDTA-Free (Roche # 11873580001) and Phosphatase Inhibitor Cocktail Set II (Millipore #524625) for 30 minutes at 4°C. Cell debris was precipitated by centrifugation at 21130 rcf for 20 minutes at 4°C, and 25μL of cell lysate was saved for input samples. Lysates were then pre-cleared using 25 μL of Pierce Protein A agarose beads (Thermo Fisher Scientific #20333). ChromoTek V5-Trap Magnetic Agarose beads (Proteintech #v5tma) and ChromoTek DYKDDDDK Fab-Trap™ Agarose (Proteintech #ffa) were used for immunoprecipitation. Precleared lysates were incubated with 25 μL of either V5 or Fab-Trap beads for 1 hour. After binding beads were washed with dilution buffer (10mM Tris-Cl pH 7.5, 150mM NaCl, 0.5mM EDTA pH 8.0) for 3 times (15 minutes each). Protein samples were resolved on SDS PAGE gels (BioRad 456-1086) along with Precision Plus Dual Color Protein Standard (Biorad 1610394) and transferred to a nitrocellulose membrane (Biorad 170-4270) using a Trans-Blot Turbo machine (Bio-Rad). Membrane was blocked with Intercept Blocking Buffer (Li-Cor #927-70001) for 1 hour at room temperature and then incubated with primary antibodies mouse anti-V5 (Invitrogen R960-25; 1:5000), rabbit anti-FLAG (Sigma-Aldrich F3165; 1:2,000), and Mouse anti-GAPDH (1:20,000; Proteintech # 60004-1-IG) overnight at 4°C. Blots were washed four times (10 minutes each) with PBT (1xPBS + 0.01% Tween-20), then incubated with secondary antibodies anti-mouse IgG-800 (LI-COR Biosciences, 1:20,000) and anti-rabbit IgG-680 (LI-COR Biosciences, 1:20,000) for 1 hour at room temperature. Blots were rinsed with PBT four times (10 minutes each) and then imaged using a Li-Cor Odyssey CLX machine. For analysis of Ds in wing discs, discs from third-instar larvae were dissected in Ringer’s Solution supplemented with Protease Inhibitor Cocktail and Phosphatase Inhibitor Cocktail on ice and then lysed in 2x Laemmli Sample Buffer (Biorad #1610737EDU) and heat denatured at 95*°*C for 5 minutes.

### Statistical analysis

Statistical analysis was performed using GraphPad Prism 10. Statistical tests for graphs showing wing area, roundness, and relative cross-vein distance were determined by Paired t-tests. All quantifications for adult wing images are presented as the mean ± SD. Quantifications for western blots are presented as the mean ± SEM. For all statistical tests, ns indicates P>0.05, * indicates P≤0.05, ** indicates P≤0.01, *** indicates P≤0.001, **** indicates P≤0.0001.

## Acknowledgments

This research used antibodies obtained from the Developmental Studies Hybridoma Bank, fly stocks from the Bloomington *Drosophila* Stock Center, plasmid from *Drosophila* Genomics Resource Center, information from Flybase, and microscopes at the Waksman Institute Shared Imaging Facility. We thank K. Mansuria for assistance with plasmid midipreps, and T. Kwok and S. Gidugu for assistance with adult wing image analysis. This research was supported by National Institutes of Health grant GM131748 (KDI).

## Notes

### Competing Interest Statement

The authors have declared no competing interest.

